# Non-apoptotic caspase-dependent regulation of enteroblast quiescence in *Drosophila*

**DOI:** 10.1101/707380

**Authors:** Lewis Arthurton, Dominik Antoni Nahotko, Jana Alonso, Luis Alberto Baena-Lopez

**Author notes:** Authors with equal contribution (Dominik Antoni Nahotko, Jana Alonso).

## Abstract

Caspase malfunction in stem cells often instigates the appearance and progression of multiple types of cancer, including human colorectal cancer. However, the caspase-dependent regulation of intestinal stem cell properties remains poorly understood. Here, we demonstrate that *Dronc*, the *Drosophila* ortholog of *caspase*-9/2 in mammals, limits the proliferation of intestinal progenitor cells and prevents the premature differentiation of enteroblasts into enterocytes. Strikingly, these unexpected roles of *Dronc* are non-apoptotic and have been uncovered under experimental conditions without basal epithelial turnover. A novel set of genetic tools have also allowed us to correlate these *Dronc* functions with its specific accumulation and transient activation in enteroblasts. Finally, we establish that the *Dronc*-dependent regulation of enteroblast quiescence, largely relies on the fine-tuning of the Notch and Insulin-TOR signalling pathways. Together, this data provides novel insights into the caspase-dependent but non-apoptotic modulation of enteroblast differentiation in non-regenerative conditions. These findings could improve our understanding regarding the origin of caspase-related intestinal malignancies, and the efficacy of therapeutic interventions based on caspase-modulating molecules.

## INTRODUCTION

Caspases are cysteine-dependent aspartate-specific proteases commonly associated with the implementation of apoptosis [1]. Despite this canonical function, an emerging body of evidence is also attributing a non-apoptotic and regulatory role to these enzymes in a wide variety of cell types, including stem cells [2]. However, the caspase-dependent control of intestinal stem cell properties and their contribution to human colon cancers remain poorly characterised. Equally unclear are the molecular mechanisms controlling the differentiation of the intermediate intestinal precursors referred to as enteroblasts (EBs), in experimental situations without associated tissue-damage and regeneration. Addressing these questions could improve our understanding regarding the origin and progression of intestinal tumours associated with caspase malfunction.

The evolutionary conservation of gene function and ease of gene manipulation in *Drosophila melanogaster* have been routinely exploited to uncover many genetic networks and cellular processes connected with human diseases [3]. Accordingly, important discoveries regarding intestinal stem cell biology and caspases have been obtained using this model organism [4]. The caspases are expressed as pro-enzymes that only become fully active after one or more steps of proteolytic processing [1, 2, 5–8]. In *Drosophila*, the apoptosis programme is initiated by the upregulation of different pro-apoptotic proteins (Hid, Reaper, Grim and/or Sickle) [7–10], which counteract molecular effects of the *Drosophila* inhibitors of apoptosis, DIAP-1 [11, 12] and -2 [13, 14]. In pro-apoptotic conditions, the main *Drosophila* initiator caspase, referred to as Dronc (*Death regulation Nedd2-like* caspase; *caspase*-2/9 orthologue in mammals) can interact molecularly with Dark-1 (Apaf-1) forming a protein complex termed the apoptosome. These events facilitate the full activation of *Dronc* [15–18], which subsequently leads to the cleavage of the effector caspases (*Death caspase-1, DCP-1* (*caspase*-7); the *death related ICE-like caspase, drICE* (*caspase*-3); *Death associated molecule related to Mch2 caspase*, *Damm* and the *Death executioner caspase related to Apopain/Yama, Decay*). Upon cleavage, functional effector caspases disrupt all of the essential subcellular structures leading to cell death [2, 6]. Intriguingly, in a previous report we uncovered a stereotyped pattern of non-apoptotic caspase activation in the *Drosophila* intestine of unknown origin and functional relevance [19].

The *Drosophila* intestine comprises of a subset of intestinal stem cells (ISCs), responsible for the renewal of the epithelial intestine [20–22]. ISCs can also differentiate upon demand as either intermediate progenitor cells termed enteroblasts (EBs) or fully differentiated secretory cells called enteroendocrine cells (EEs) [23]. The EBs rarely, if ever divide but can terminally differentiate as mature absorptive cells referred to as enterocytes (ECs) [23]. Throughout the last two decades, an abundant body of literature has emerged describing many of the genetic factors controlling the proliferation and differentiation of ISCs into EBs; however, the differentiation pathway of EBs to ECs remains less well characterised. Notch signalling is one of the instrumental signalling cascades permitting the entry of the EBs into the EC differentiation programme. The interaction of the extracellular domain of the Notch receptor with its ligands (either Delta or Serrate) facilitates the activation of this evolutionary conserved signalling cascade [24]. Notch activation culminates with the release and subsequent translocation into the nucleus of the Notch intra-cellular domain (Notch^Intra^) [25, 26]. The interaction of the Notch^intra^ fragment with several transcription factors governs the transcriptional response in a highly cell-specific manner [27]. Low levels of Notch-signalling promote the self-renewal of ISCs, whilst elevated Notch activation stimulates the conversion of ISCs into EB [28]. In addition to other transcriptional effects, high levels of Notch in EBs repress the expression of the tuberous sclerosis protein complex 1 and 2 (TSC-1 and TSC-2) [29]. These proteins are negative regulators of the Insulin-TOR pathway, and therefore naturally limit cellular growth [29]. Conversely, insulin-TOR pathway upregulation in response to Notch activation instigates the entry of EBs into the EC differentiation programme [29, 30]. In tissue damaging conditions, Notch activation can also regulate additional signalling pathways leading to differentiation activity in EBs [31]. Interestingly, caspase malfunction has been shown to alter the proliferation and differentiation of ISCs in regenerative conditions [32]. Furthermore, *caspase*-9 deficiency in patients suffering from colorectal cancer appears to stimulate the proliferation of intestinal precursors, whilst compromising their differentiation [33]. In this manuscript we describe how the specific protein accumulation and non-apoptotic activation of *Dronc* promotes EB quiescence by preventing their premature entry into the EC differentiation pathway. Importantly, these novel *Dronc* functions are accomplished without the participation of effector caspases. Alternatively, they seem to require the *Dronc*-dependent regulation of Notch signalling and Insulin-TOR pathways.

## RESULTS

### Robust non-apoptotic caspase activation pattern in the Drosophila intestine independent of cellular turnover

Using a highly-sensitive caspase activity sensor (Drice-based-sensor-QF; DBS-S-QF), we previously reported the presence of a stereotyped pattern of non-apoptotic caspase activation in the adult posterior midgut of *Drosophila* (Fig 1A) [19]. Following this initial observation, we sought to investigate the potential correlation of this caspase activation with intestinal homeostasis at the cellular level, monitoring the dynamics of cell proliferation and differentiation. To that end, we utilised the ReDDM cell lineage tracing system [34]. This system employs the combined expression of a short-lived GFP-marker and a semi-permanent Histone-RFP-labelling to readily visualise the turnover of the intestinal cells [34]. The short-lived GFP labelling co-exists with the Histone-RFP marker within all undifferentiated intestinal progenitor cells (ISCs and EBs), expressing the Gal4 protein under the regulation of the *esg* promoter (*esg*-expressing cells, Fig 1B) [34]. However, the silencing of the *esg* promoter during differentiation stimulates the rapid degradation of the GFP signal whilst the Histone-RFP remains. The stability of the Histone-RFP bound to the DNA, and the absence of GFP signal, allows the identification of differentiated ECs and EEs after the Gal4 protein production ceases [34]. The incorporation of a Gal80 thermosensitive repressor of the Gal4 protein, under the regulation of the *Tubulin* promoter (TubG80^ts^) facilitates the spatial and temporal control of the ReDDM system [34]. We exploited this dual labelling system to distinguish undifferentiated intestinal progenitors (expressing GFP and RFP) from their progeny (Histone-RFP-positive cells) in experimental conditions with and without epithelial replenishment. The REDDM cell lineage-tracing analysis of adult flies reared in our experimental conditions (see fly food composition in materials and methods and the experimental regime in Appendix Fig 1A) failed to show signs of regeneration during the first 7 days post ReDDM induction (Fig 1A-C); note the almost perfect overlap between the red and green fluorescent signals within intestinal precursors (*esg*-positive cells) indicates the absence of differentiation [34] (Fig 1B-C). By contrast, unspecific tissue damage triggered by either exposure to paraquat (an organic compound that induces the production of reactive oxygen species [35]) or detrimental dietary conditions, induced robust labelling of the intestines with the DBS-S-QF caspase sensor, and increased the number of differentiated ECs (Histone-RFP alone; compare Fig 1B-C with either Fig 1E-F or Appendix Fig 1C-D). These results indicated the ability of our experimental conditions to preserve the intestinal epithelia in a quiescent state, without homeostatic cellular death and turnover during the first 7-days post ReDDM activation. To further consolidate this conclusion, we overexpressed either one or two copies of the effector-caspase inhibitor P35 [36], in intestinal progenitor cells. This genetic manipulation did not increase the number of either progenitor or differentiated cells in our experimental conditions, as one would expect if apoptosis and cell replenishment would be taking place (Fig 1G-I). Furthermore, P35-expressing intestines were cellularly and morphologically equivalent to the non-P35 expressing controls (compare Fig 1B and 1H). These results unambiguously confirmed the quiescent status of the intestines maintained under our experimental regime, and the non-apoptotic nature of the caspase activation detected with the DBS-S-QF reporter [19]. Paradoxically, it also raised the question of whether there was a functional requirement for the described non-apoptotic pattern of caspase activation, since the epithelial integrity of the gut was unaffected by the overexpression of P35.

**Figure 1:**
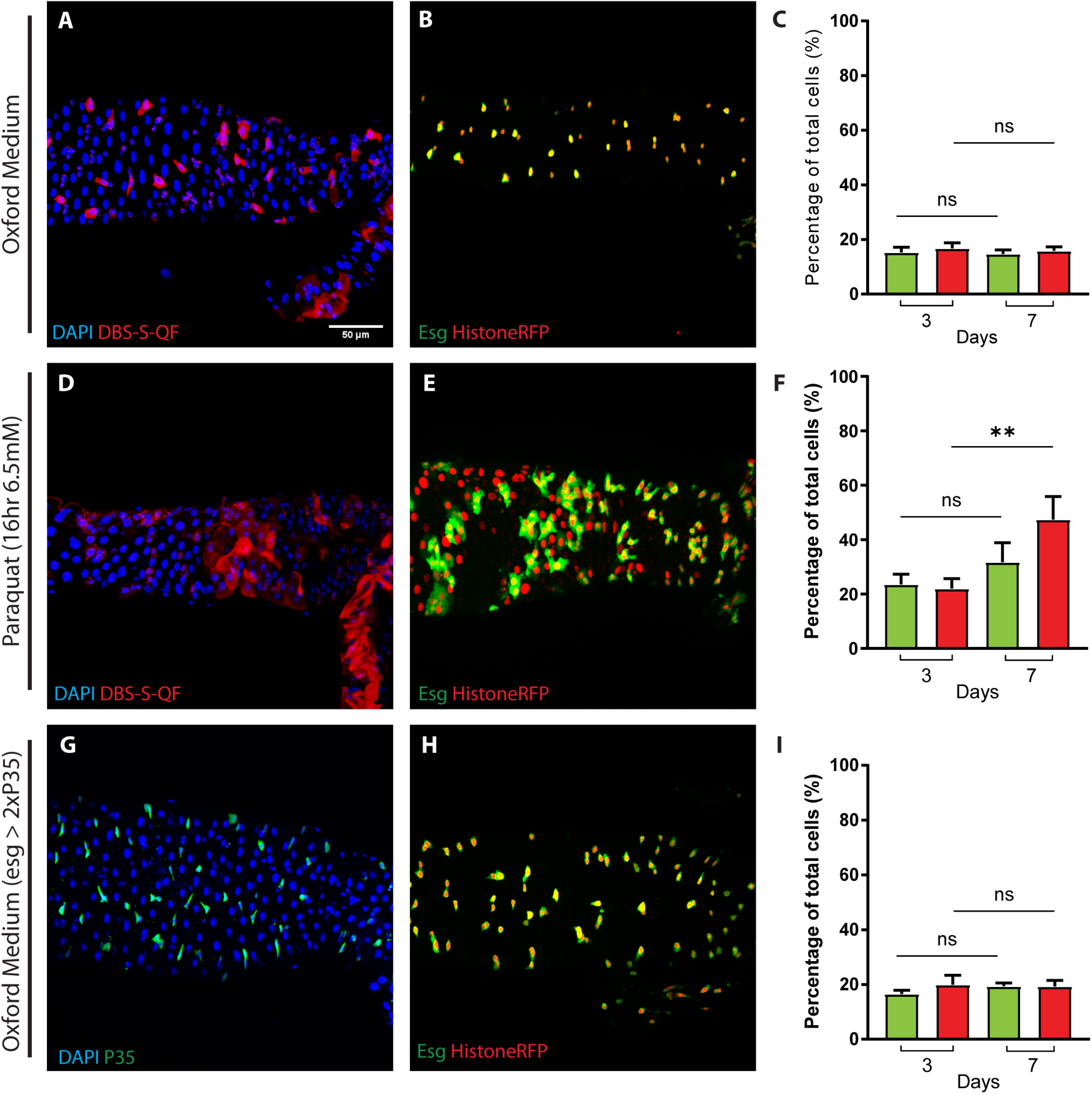
Non-apoptotic caspase activation independent of basal cellular turnover in the *Drosophila* Intestinal System. (A) Adult female posterior Intestine (Region 4-Region 5) of *Drosophila* reared on Oxford Medium activating the initiator caspase reporter DBS-S-QF (Red, immunostaining anti-HA). (B) Representative image of ReDDM activation in a *Drosophila* intestine reared in Oxford Medium and following an experimental regime that protects the epithelial integrity (transferring flies every two days into vials with fresh food). *esg* (green fluorescent signal) labels intestinal progenitor cells, whilst Histone-RFP (red fluorescent signal) acts as a semi-permanent labelling of differentiating cells (B). Notice the extensive overlap between the two markers in *esg*-expressing cells and therefore the absence of differentiated cells only showing the Histone-RFP labelling (B), as an indication of negligible epithelial turnover (B). (C) Quantification of intestinal cell subpopulations labelled with ReDDM system (GFP and Histone-RFP) in Oxford Medium and experimental conditions that protect epithelial integrity at different time points post ReDDM activation (3- and 7-days); note that none of the cell populations in the gut (GFP (P = 0.5267) or Histone-RFP (P=0.2752)) significantly increase in number overtime (Mann-Whitney, 3d N=34, 7d N=45; C). (D) Intestinal DBS-S-QF reporter activation (red, anti-HA immunostaining) observed in Oxford Medium after 16 hours of paraquat treatment; note the expansion of the labelling with DBS-S-QF to large intestinal cells (ECs) in this case (compare D with A). (E) Representative example of the ReDDM lineage-tracing in a *Drosophila* intestine reared in Oxford Medium and paraquat (20 mM) over 16 hours; notice the abundance of Histone-RFP cells without GFP signal, as an indication of epithelial damage and subsequent differentiation of progenitor cells. (F) Quantification of ReDDM labelling after paraquat treatment; notice the statistically significant increase (P = 0.0099) of Histone-RFP expressing cells without GFP signal (Unpaired two-tailed t test, 3d n = 9, 7d N = 7). (G) Representative example of an intestine expressing two copies of the effector caspase inhibitor P35 under the regulation of *esg*-gal4 (2x P35, green immunostaining with antibody against P35). (H) ReDDM lineage-tracing system in a *Drosophila* intestine, reared in Oxford Medium and protective experimental conditions which protect epithelial integrity, expressing two copies of the effector caspase inhibitor P35 under the regulation of *esg*-Gal4. (I) ReDDM quantification corresponding to the intestines described in (H); no significant increase in either *esg* (P = 0.1352) or Histone-RFP (P = 0.9801) cell number is observed (unpaired two-tailed t test, 3d N = 12, 7d N = 11). Error bars represent Standard Error of the Mean in all panels. In the histograms depicted in C, F and I, green bars correspond to GFP and red bars correspond to HistoneRFP. DAPI (blue) labels the nuclei in panels A, D and G.

### Dronc prevents the premature differentiation of intestinal progenitor cells

The lack of cellular and morphological phenotypes linked to the overexpression of P35 could suggest a negligible functional requirement for the non-apoptotic caspase activation previously described. However, P35 overexpression only prevents the activity of effector caspases, and therefore a potential function of the initiator caspases could have been overlooked. To address this question, we created a new *Dronc* knockout allele using genome engineering protocols [37]. This resulted in a *Dronc* allele (*Dronc*^KO^) which contained an attP integration site immediately after the *Dronc* promoter and within the 5’UTR of the gene (Appendix Fig 2A and B). As with previously described *Dronc* null alleles [17, 38], the new mutant was homozygous lethal during early pupal development, and it failed to genetically complement other *Dronc* mutations (Appendix Fig 2C). The heterozygous insertion of a wild-type *Dronc* cDNA into the *Dronc* attP-site gave rise to fertile adult flies that appeared largely similar to their wild-type siblings (Appendix Fig 2D-F). These results indicated that our rescue construct retained all the essential functionality of the endogenous gene, whilst validating the attP integration site. Next we created a conditional allele of *Dronc* (*Dronc*^KO-FRT Dronc-GFP-Apex FRT-QF^; Appendix Fig 2G). This allele contained functional cDNA of *Dronc* flanked by FRT recombination sequences which is able to rescue various *Dronc* null mutant alleles. The excision of the FRT-rescue cassette enables the efficient elimination of *Dronc* expression in any cell type of interest, including those with low rates of proliferation such as the EBs. Additionally, we placed in this allele the sequence for the QF transcriptional activator downstream of the FRT-rescue-cassette (*Dronc*^KO-FRT^ ^Dronc-GFP-Apex^ ^FRT-QF^). Since the QF factor can induce the expression of any cellular marker of interest upon binding to the QUAS sequences (e.g QUAS-LacZ), this feature can be used to identify FRT-cassette excision events in all of the cells physiologically transcribing the QF protein under the regulation of the *Dronc* promoter. To determine the excision efficacy of our allele and potential loss-of-function (LOF) phenotypes linked to *Dronc* in the intestinal progenitor cells, we induced the expression of a Flippase recombinase using the *esg*-Gal4 driver in a *Dronc* heterozygous mutant background. 3 days after Flippase induction and FRT-cassette excision, 91.35% of *esg*-labelled cells (GFP-positive cells) showed transcriptional activation of the *lacZ* reporter gene (Appendix Fig 3A-B). These results indicated the suitability of our allele to assess the role of *Dronc* within intestinal progenitor cells. Additionally, we noticed signs of tissue differentiation and regeneration (i.e increased number of enlarged LacZ-positive cells without *esg* expression (GFP-negative cells); Appendix Fig 3A). Equivalent results were obtained using a different conditional allele that expressed a Suntag-HA-Cherry chimeric protein upon the FRT-rescue cassette excision (*Dronc*^KO-FRT Dronc-GFP-Apex FRT-Suntag-HA-Cherry^) (Fig 2B-E; Appendix Fig 2H). Furthermore, we observed hyperplasia (Fig 2C), cellular and nuclear enlargement (Fig 2D and Appendix Fig 3C), co-expression of Histone-RFP with the EC maker Pdm-1, and the co-localisation between the GFP and Pdm1 (inset in Fig 2B and 2E and Appendix Fig 3D). These phenotypes also worsened over time (Fig 2C-E). To discard any unspecific/detrimental effect linked to the Suntag-HA-cherry peptide, flies expressing a WT version of this caspase member tagged with Suntag-HA-Cherry (*Dronc*^KO-FRT^ ^Dronc-GFP-Apex FRT-DroncWT-Suntag-HA-Cherry^) failed to show any of the previously described phenotypes (Appendix Fig 2I and 3E). Collectively, these results suggested that *Dronc* insufficiency in the intestinal progenitor cells causes gut hyperplasia and the premature entry in the EC differentiation program. Since our data was collected in experimental conditions without cellular turnover, *Dronc* seemed to restrain EBs in a quiescent state. To further characterise the differentiation features of *Dronc* mutant progenitor cells, we next analysed the expression profile of metabolic enzymes highly enriched in ECs [39]. Interestingly, we found that several of these genes were transcriptionally downregulated in our mutant conditions, while others were upregulated (Appendix Fig 3F-H). This irregular expression of EC markers confirmed the premature and/or defective differentiation status of *Dronc*-mutant progenitor cells.

**Figure 2:**
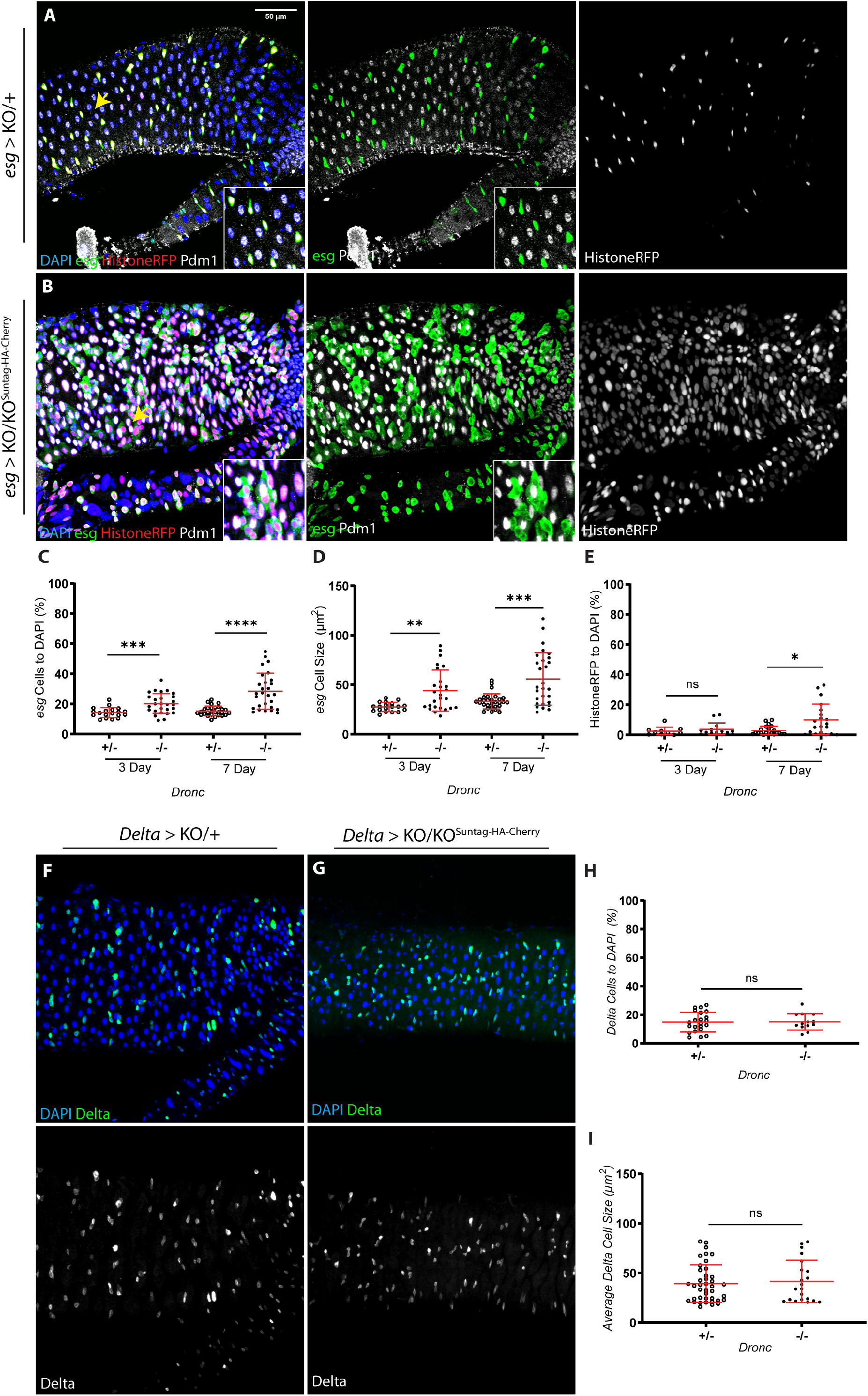
*Dronc* facilitates Enteroblast quiescence in experimental conditions without basal cellular turnover. Representative image of ReDDM activation 7 days (7d) after temperature shift at 29 C°, *Dronc* heterozygous intestine, reared in Oxford Medium and an experimental regime which protects epithelial integrity; *esg* expression (green) labels the intestinal progenitor cells, Histone-RFP (red) is a semi-permanent marker retained in differentiated cells and Pdm-1 (grey) labels differentiated ECs. *Dronc* KO progenitor cells show EC features; increased *esg* cellular size and premature expression of the EC marker Pdm-1 (compare panels and detailed insets from A with B). (C) Relative number of *esg*-expressing cells normalised to DAPI; notice that the relative percentage *of esg*-labelled cells is significantly higher in *Dronc* KO mutant conditions at 3 days (P = 0.0007) and 7d (p= <0.0001) post temperature shift at 29 C° (unpaired two-tailed t-test, +/− N=32, −/− N=25). (D) Average cell size of *esg*-expressing cells (μm^2^); notice the increased cell size of *Dronc* KO progenitor cells at 3d (P = 0.0014) and 7d (P = <0.0001) post temperature shift at 29 C° (unpaired two-tailed parametric t-test, 3d +/− N = 19, 3d −/− N = 28, 7d +/− N = 33, 7d −/− N = 28). (E) Relative number of *esg*-negative cells expressing Histone-RFP normalised to DAPI; notice that the number of Histone-RFP cells without *esg* expression is significantly higher in *Dronc* KO mutant conditions at 7d (P = 0.0046) (Mann-Whitney test, +/− N = 24, −/− N = 20). (F) Representative image of a 7d *Dronc* heterozygous intestine in which intestinal stem cells express GFP under the regulation of *Dl*-Gal4. (G) Representative example of a 7d post temperature shift at 29 C° Intestinal Stem Cell *Dronc* KO homozygous mutant intestine in which intestinal stem cells express GFP under the regulation of *Dl*-Gal4; there are no morphological differences between F and G. (H) Relative number of *Delta*-expressing cells normalised to DAPI; there is no significant increase in intestinal stem cell number between heterozygous and homozygous *Dronc* mutant conditions (P = 0.9231, unpaired two-tailed t test, +/− N = 22, −/− N = 13). (I) Average *Delta* cell size (μm^2^); notice that the cell size does not change between heterozygous and homozygous *Dronc* mutant ISCs (P = 0.9694, unpaired two-tailed t test, +/− N = 42, −/− N = 21). Error bars represent Standard Deviation of the Mean in all panels. DAPI (blue) labels cell nuclei in panels A, B, F and G. All of the experiments described in the figure were performed in Oxford Medium following an experimental regime that protects the epithelial integrity.

### The Dronc-dependent quiescent-state of intestinal progenitor cells requires its enzymatic activity but does not involve the effector caspases

Previous literature has demonstrated the ability for Dronc to regulate signalling events independently of its enzymatic activity, through protein-protein interactions [40]. Following the demonstration of the non-apoptotic function of *Dronc* in progenitor cells, we investigated whether the catalytic activity or merely the presence of the protein was required for these functions. To distinguish between these two possibilities, we utilised a conditional allele of *Dronc* that can express an enzymatically inactive form (containing C318A and E352A amino acid substitutions) of the protein (FL-CAEA; *Dronc*^KO-FRT Dronc-GFP-Apex FRT-Dronc FL-CAEA-Suntag-HA-Cherry^; Appendix Fig 2J) upon Flippase mediated-excision of the upstream FRT-rescue cassette. The expression of this mutant protein in progenitor cells caused less penetrant phenotypes in terms of proliferation, but an equivalent phenotype from a differentiation perspective (cell size increase and expression of EC differentiation markers; Appendix Fig 3I-N). We further validated these observations by expressing a catalytically inactive version of *Dronc* in which the CARD domain (a protein-protein interaction domain) was also deleted (dCAEA; *Dronc*^KO-FRT Dronc-GFP-Apex FRT-Dronc dCAEA-Suntag-HA-Cherry^; Appendix Fig 2K and 3l-N). These findings strongly suggested the molecular association of the Dronc functions with its catalytic activity. Next, we explored whether these functions could be performed by the primary substrates of Dronc in many cellular contexts, the effector caspases (*drIce, Dcp-1, Decay* and *Damm*). Since the overexpression of two copies of P35 did not alter the cellular or morphological features of the gut, our previous experiments already discarded a potential enzymatic requirement of effector caspases. However, a potential role of these proteins acting as scaffolding partners could still exist. To investigate this possibility, we simultaneously targeted the expression of all of these downstream caspase members using validated RNAi lines [41]. This experimental design prevents the previously described functional redundancy between the effector caspases [42]. This set of experiments failed to replicate the excess of proliferation and premature differentiation phenotypes observed in *Dronc* LOF conditions (Appendix Fig 4). Together, these results strongly argue in favour of a *Dronc* specific regulation of progenitor cell properties through its enzymatic activity, but fully independent of effector caspases and the apoptosis programme (see discussion).

### The non-apoptotic function of Dronc is exclusively required in EBs

Although our previous experiments uncovered unknown functions of *Dronc*, they could not discriminate whether these functions were ascribed to the ISCs, the EBs or both cell types. To address this question, we specifically targeted the expression of *Dronc* in ISCs using the *Delta*-Gal4 driver [43]. As previously shown with *esg*-Gal4, we obtained a high excision efficiency (81.15%) of the FRT-rescue cassette in ISCs by driving the expression of the Flippase recombinase with the *Delta*-Gal4 line (Appendix Fig 3O-P). However, *Dronc* deficiency in ISCs did not cause any cellular or morphological alteration of the gut, and no-increase in the number and cell size of Delta-positive cells (Fig 2H and I). To explore the possibility of increased differentiation into EE fate following *Dronc* LOF, we quantified the number of small nuclei concomitantly the EE cell identity marker, Prospero. No statistical differences were observed in this set of experiments between experimental and control intestines (Appendix Fig 3Q). These results unambiguously indicated a specific functional requirement for *Dronc* in EBs. They also explained the differentiation bias of progenitor cells towards EC fate utilising the *esg-Gal4* driver (see discussion).

### The protein accumulation and transient activation of Dronc in EBs determines its functional specificity

Our previous data suggested a functional requirement for *Dronc* in EBs, but the molecular origin of such specificity remained elusive. To elucidate this question, we first explored whether the transcriptional regulation of *Dronc* could be restricted to EBs, using a newly created *Dronc*^KO-Gal4^ strain (Appendix Fig 2L). This fly line transcriptionally expresses Gal4 under the physiological regulation of the *Dronc* promoter, and therefore is a *bona fide* transcriptional read out of the gene. *Dronc* was widely transcribed in all of the intestinal cell subtypes (Appendix Fig 5A). This experiment separated the specificity of the *Dronc*-dependent EB quiescence from the transcriptional regulation of the gene. Next, we investigated whether the *Dronc* protein level could be differentially regulated in EBs. Our laboratory has created multiple *Dronc* alleles tagged at the C-terminal with different peptides (e.g HA, Cherry and GFP) that successfully rescue the insufficiency of *Dronc* amorphic alleles. However, we failed to detect the physiological expression of Dronc using these fly strains in most of the analysed tissues, including the intestinal system. Similar frustrating results were obtained using the short repertoire of validated antibodies raised against the Dronc protein [44, 45]. To circumvent this technical issue, we created a new Dronc protein sensor. The new reporter line ubiquitously expresses a catalytically inactive (C318A) and tagged template of *Dronc* under the regulation of the *Actin* promoter. This Dronc mutant cannot induce apoptosis, however the tagging at the C-terminus with a modified GFP and Myc facilitates its immunodetection (Appendix Fig 5B). Importantly, the overexpression of this construct generates fertile adult flies without any noticeable developmental or morphological defects. Furthermore, this reporter was able to recapitulate the subcellular localisation of Dronc previously described in other tissues such as the salivary glands and the wing imaginal discs (Appendix Fig 5C) [46]. Interestingly, our construct reported a noticeable accumulation of Dronc protein levels within a subpopulation of progenitor cells despite the absence of cellular turnover in our experimental conditions (Fig 3A). Furthermore, Dronc accumulation showed a striking overlap with the EB marker Su(H) (Fig 3B). Complementing these findings, we also observed that the specific EB accumulation of Dronc disappeared in tissue-damaging conditions, after exposure to paraquat (Appendix Fig 5D). Considering these findings, we decided to re-evaluate our previous results regarding the activation of the DBS-S-QF sensor, combining this activity reporter line with Su(H). These experiments demonstrated that most of the caspase activation in intestinal progenitor cells was specifically ascribed to the EBs in our experimental conditions (Fig 3C and Appendix Fig 5E). These results strongly correlated the non-apoptotic functions of *Dronc* with its specific accumulation and transient activation in EBs.

**Figure 3:**
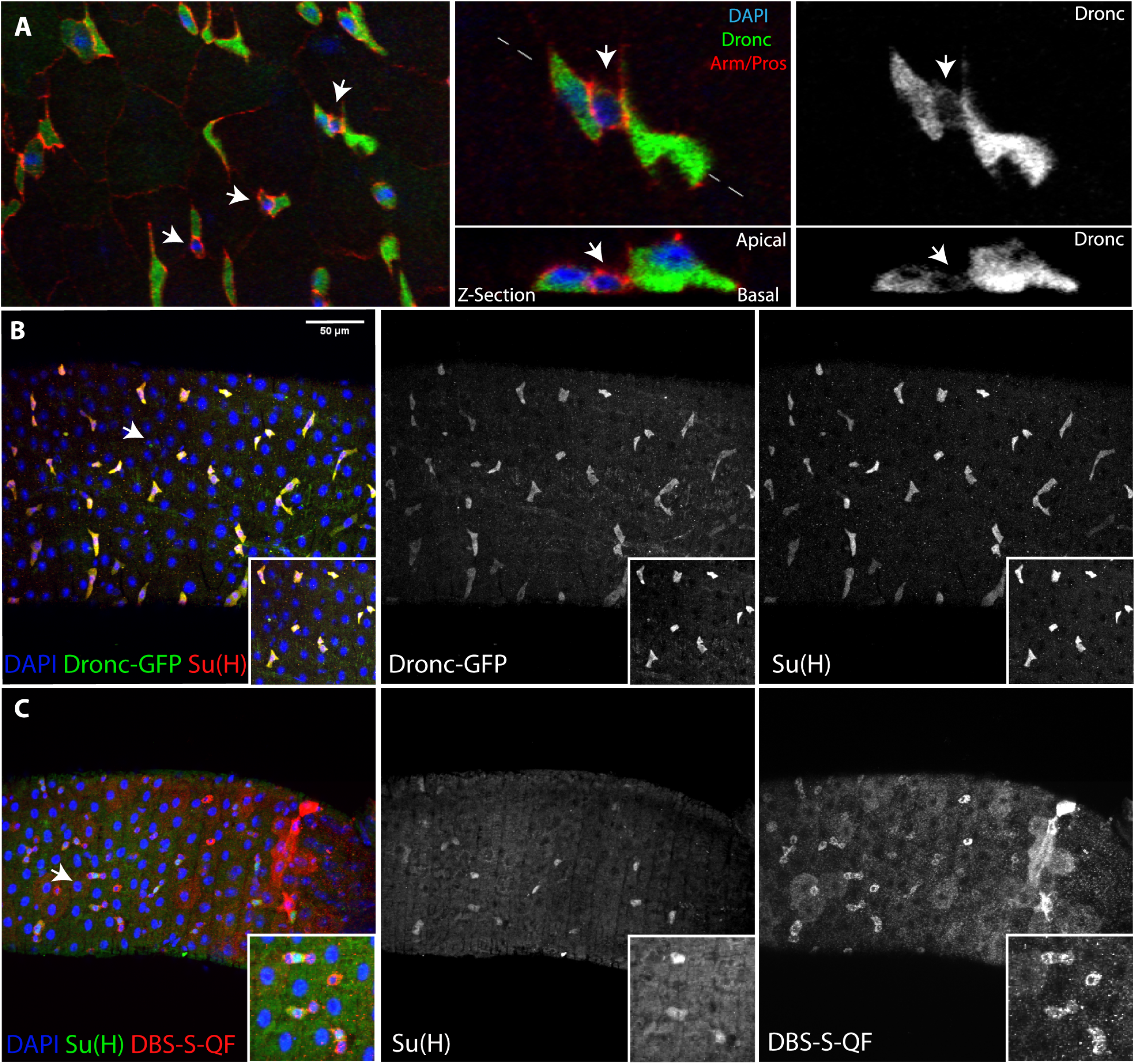
Dronc is preferentially accumulated and activated in quiescent enteroblasts. (A) Representative image of a *Drosophila* posterior midgut showing the expression of Armadillo (red membranes), Prospero (red nuclei of EEs) and Dronc (green, inmmunostaining against GFP); notice that Dronc is preferentially enriched in a subpopulation of intestinal progenitor cells (white arrows indicate intestinal progenitor cells showing lower levels of Dronc). (B) The enteroblast marker Su(H) (red, immunostaining against Beta-galactosidase) strongly co-localises with high levels of Dronc expression (green, immunostaining against GFP); white arrow indicates the enlarged area depicted in the insets (B). (C) There is extensive overlap between the expression of the EB marker Su(H) (green, immunostaining against Beta-galactosidase) and the apical caspase reporter DBS-S-QF (red, immunostaining against HA). DAPI (Blue) labels cell nuclei in all the panels (A-C). All of the experiments described in the figure were performed in Oxford Medium following an experimental regime that protects the epithelial integrity.

### Dronc regulates EB quiescence acting upstream of Notch and Insulin-TOR pathway

As stated, the Notch pathway is one of key signalling cascades involved in the regulation of cell proliferation and differentiation in the *Drosophila* and the mammalian intestinal system [28, 47, 48]. Furthermore, non-apoptotic protein-protein interactions have been described between Dronc and Notch pathway regulators (e.g. Numb) in the *Drosophila* neuroblasts [40]. Considering these precedents, we analysed through classical genetic epistasis the potential genetic interplay between Dronc and the Notch pathway. The inhibition of Notch-signalling in intestinal progenitor cells promotes the expansion of ISCs and EEs, whilst preventing the differentiation of EBs to ECs [49]. Since *Dronc* LOF facilitates the premature differentiation of EBs, we first investigated whether Notch-signalling deficiency would be able to revert the *Dronc* mutant phenotype. To that end, we simultaneously targeted the expression of *Dronc* and *Notch* in progenitor cells using the *esg*-Gal4 driver. The epithelial features of intestines obtained from these experiments were equivalent to the Notch LOF conditions, indicating that the characteristic increase in cell size and EC features linked to the *Dronc* insufficiency were supressed (compare Fig 4A-C with Fig 2B-E, Appendix Fig 3A and I-N). However, we did detect a mild increase in cell number in the *Dronc* LOF intestines, perhaps linked to a potential rescue of apoptosis (Figure 4C). Conversely, the ectopic activation of Notch-pathway in *Dronc* mutant progenitor cells promoted their quick conversion into ECs and subsequent elimination from the epithelia (Fig 4D-E). Furthermore, the depletion of intestinal precursors was quicker than with the activation of Notch (N^intra^) in the control genetic background (Fig 4F). These results genetically located the function of Dronc upstream of the Notch-pathway, likely acting as a negative regulator of the terminal differentiation programme of EBs to ECs. The Insulin-TOR pathway is required downstream of the Notch-pathway to complete the terminal differentiation of EBs [29, 30]. Since our data suggested that *Dronc* LOF could boost Notch-signalling and ultimately differentiation, we formally tested whether the Insulin-TOR pathway would be genetically downstream of the *Dronc* and Notch-pathway. To that end, we concomitantly eliminated the expression of *Dronc* and the Insulin receptor. As expected, the lack of Insulin-TOR signalling rescued the premature differentiation phenotypes triggered by the Dronc LOF (Fig 4G-I). Collectively, these results confirm the genetic hierarchy established between Notch and Insulin-TOR signalling pathways during the terminal conversion of EBs to ECs [29, 30], whilst placing *Dronc* as an upstream regulator of Notch-pathway.

**Figure 4:**
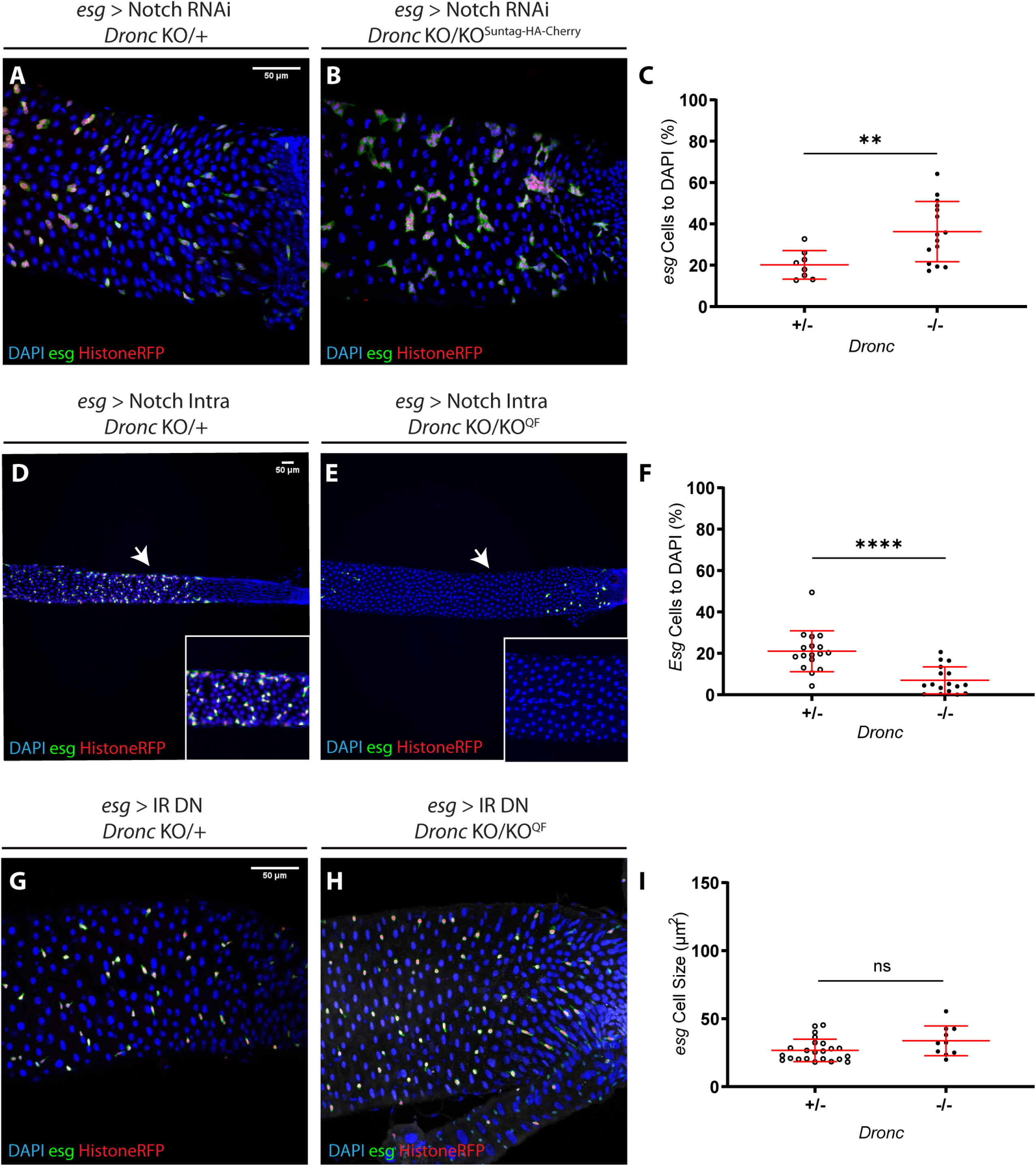
The *Dronc*-dependent EB quiescence demands the fine-tuning of Notch and Insulin pathways. Representative ReDDM labeling of a *Drosophila Dronc* heterozygous intestine overexpressing an RNAi against Notch for 3 days; notice the lack of fully differentiated Histone-RFP cells as EC (red) without *esg* expression (green, GFP). (B) Representative ReDDM labeling of a *Drosophila Dronc* mutant homozygous intestine overexpressing an RNAi against Notch for 3 post temperature shift at 29 C°; notice the lack of differentiated Histone-RFP cells as EC (red) without *esg* expression (green, GFP), as well as the increase of GFP-positive cells (compare A to B). (C) Quantification of the number of Histone-RFP cells normalised to DAPI (proxy of progenitor cell proliferation obtained from the experiments shown in A and B panels); notice the statically significant increase in Histone-RFP positive cells in *Dronc* homozygous mutant intestines compared to controls (P =0.0080) (unpaired two-tailed t test, +/− N = 8, −/− N = 15). (D) *Drosophila Dronc* heterozygous intestine overexpressing the Notch intracellular domain for 7 days post temperature shift at 29 C°; intestinal *esg*-positive progenitor cells (green (GFP) and red (Histone-RFP). (E) *Drosophila Dronc* homozygous intestine overexpressing the Notch intracellular domain for 7d post temperature shift at 29 C°; notice that the *Dronc* deficiency accelerates the elimination of intestinal progenitor cells induced by Notch overactivation (compare D and E). The white arrows indicate the position of insets 500μm from the posterior region. Note the complete loss of *esg* labelled cells in this region. (F) Relative number of *esg*-positive cells to DAPI in either heterozygous or homozygous *Dronc* mutant *esg* cells overexpressing Notch-Intra; note the significant reduction of *esg*-expressing cells (P = < 0.0001) (Mann-Whitney test, +/− N = 17, −/− N = 17). (G) Representative ReDDM labelling of a *Drosophila Dronc* heterozygous intestine overexpressing a dominant negative form of the insulin receptor for 3d post temperature shift. (H) Representative ReDDM labeling of a *Drosophila esg Dronc* homozygous mutant intestine overexpressing a dominant negative form of the insulin receptor for 3 days post temperature shift; note that G and H show equivalent cellular morphological features. (I) Average cell size (μm^2^) of the *esg*-expressing cells corresponding to the genotypes illustrated in panels G and H; notice there is no significant change in cell size between genetic conditions (P = 0.2484, unpaired two tailed t test, +/− N = 24, −/− N = 10). DAPI staining labels cell nuclei in panels A, B, D, E, G and H. Error bars represent Standard deviation of the Mean in all panels. All the experiments described in the figure were performed in Oxford Medium with an experimental regime which protects epithelial integrity

## DISCUSSION

### The specific accumulation and activation of Dronc in EBs promotes cellular quiescence

Over the past two decades, numerous *Drosophila* studies have utilised either environmental or genetically-induced tissue damaging conditions to decipher the molecular factors controlling the proliferation and differentiation of ISCs into EBs [32, 50–52]. However, the differentiation step of EBs into fully differentiated ECs is less understood. We utilised experimental conditions, which ensure the *Drosophila* intestine remains free of apoptosis and basal cellular turnover during at least the first 7 days post ReDDM activation (Fig 1) to investigate the potential non-apoptotic role for the caspases within intestinal progenitor cells. Strikingly, in this experimental setting we observed a stereotypical pattern of caspase activation that appears to occur to a large extent in EBs (Fig. 3C, Appendix Fig 5E and [19]). Our data demonstrates the correlation of this caspase activation with the initiator caspase *Dronc*. Furthermore, we demonstrate that the accumulation and activation of Dronc in EBs is essential to prevent the appearance of gut hyperplasia, as well as the entry of these progenitor cells into the EC differentiation programme (Fig 2 and 3). These findings indicate that a sophisticated genetic network controls the differentiation of EBs, and therefore the epithelial homeostasis of the intestine does not rely exclusively on the regulation of ISCs. Based on this, the fine-tuning of EB properties could be more relevant than previously thought for maintaining the intestinal epithelial homeostasis. Importantly, *caspase*-9 deficiency in human intestinal precursors results in excessive proliferation and poor differentiation [33]. Furthermore, these features are considered a bad prognosis marker for human colon cancer [33]. Together, our findings could help to better explain the origin of these human malignancies.

Independently, our data supports the hypothesis that non-apoptotic caspase activation is key to modulate fundamental stem cell properties, such as cell proliferation and differentiation, beyond their role in apoptosis [2, 53, 54]. Indeed, considering the fast-growing list of examples indicating an implication for the caspases in non-apoptotic functions [2, 53, 54], apoptosis could be the phenotypically more apparent function of these enzymes, but not necessarily the primary and/or the most relevant. Future evolutionary analysis of these novel non-apoptotic functions in primitive organisms should clarify the primary function of caspases.

### The Dronc-dependent EB quiescence relies on its enzymatic activity of Dronc and its ability to modulate Notch signalling, but is independent of the apoptotic pathway

Our results indicate that the sole presence of Dronc is insufficient to fulfil all of its functions in EBs, and therefore its enzymatic activity is required. Therefore, Dronc does not regulate EB quiescence acting as a scaffold protein but as a proteolytic enzyme. Our experiments also suggest that either the expression or activation of effector caspases is dispensable in order to ensure EB quiescence, but instead an unknown substrate “X” of Dronc must exist (Fig 5A). Importantly, our genetic epistasis indicates that the Dronc-mediated cleavage of such a factor could directly or indirectly limit Notch-signalling. Supporting this model, we have demonstrated that *Dronc* LOF differentiation phenotypes can be rescued by supressing the signalling of either the Notch or Insulin-TOR pathways. Reciprocally, the phenotypes induced by Notch activation are enhanced in *Dronc* mutant conditions. Collectively, these findings strongly suggest a non-apoptotic function of *Dronc* as a signalling regulator in the EBs.

### Caspases can modulate intestinal cellular properties through different molecular mechanisms

An important body of literature has shown the ability of apoptotic caspase-activating cells to stimulate the proliferation and differentiation of intestinal precursors through the activation of key signalling pathways in tissue-damaging conditions [32, 50–52]. Beyond the apoptotic caspase-dependent pro-proliferative roles, a recent report has also shown that effector caspases acting downstream of Hippo-signalling mediates the cleavage of the chromatin regulator Brahma [32]. This caspase effect helps to restrain the proliferation and differentiation of ISCs after tissue damage (transition 1 in the model of Fig 5B, [32]). Our findings now demonstrate that *Dronc* is required to control the timely entry of EBs into the EC differentiation programme (transition 2 model Fig 5B). This new function of *Dronc* is unlikely to be correlated with expression of Brahma, since it is totally independent of effector caspases. However, it is tightly connected with the regulation of Notch-signalling and of the Insulin-TOR pathway (Fig 5A). Collectively, these results illustrate the ability of caspases to modulate in multiple ways the homeostasis of different subpopulations of intestinal precursors without causing apoptosis. In parallel, they suggest the presence of an unknown and highly specific mechanism to activate caspases at sublethal thresholds in different intestinal cell subpopulations.

**Figure 5:**
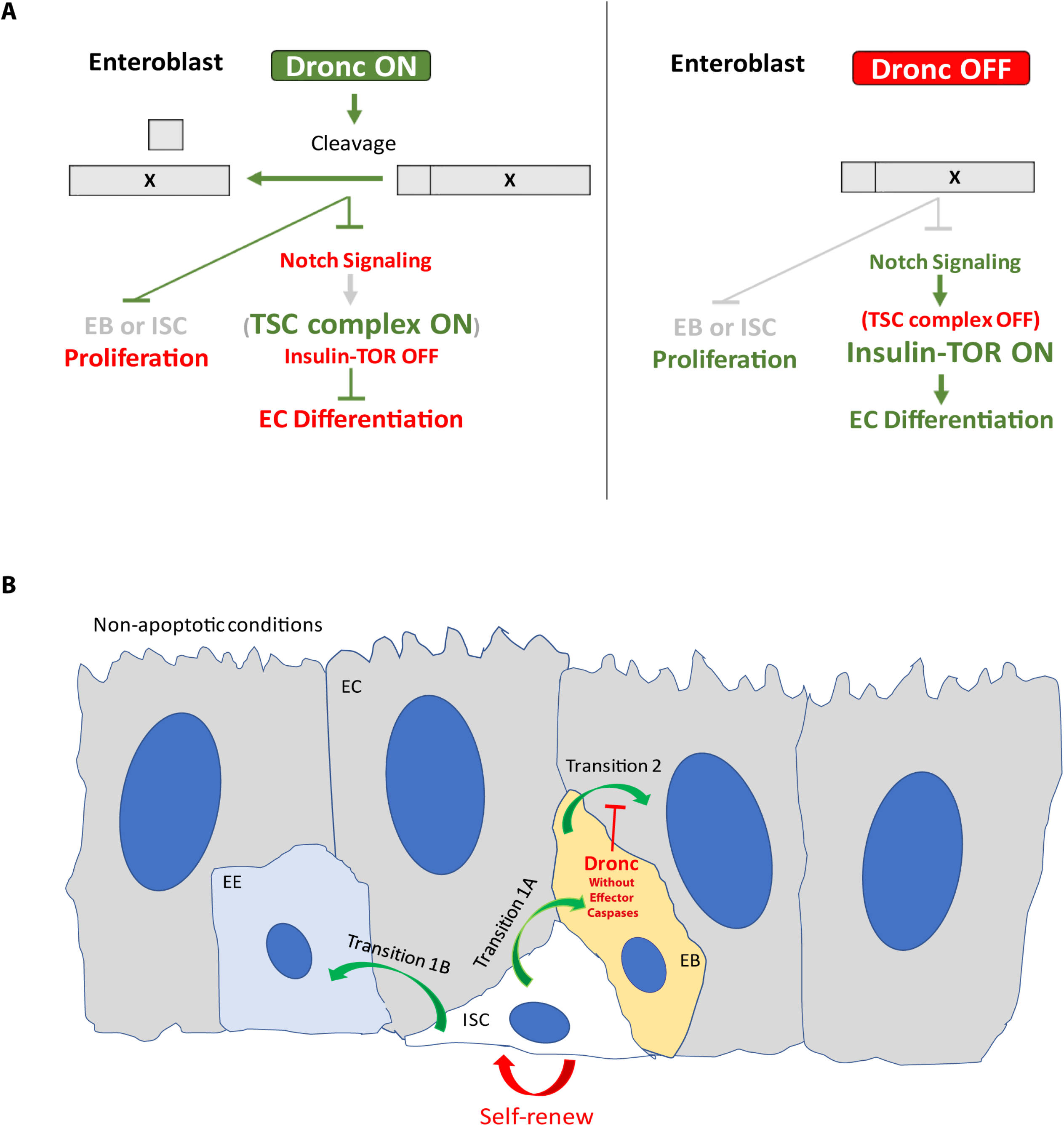
Working model summarising the interplay between *Dronc* and Notch pathway in EBs and the functional epithelial effects of *Dronc*. (A) Schematic model illustrating the non-apoptotic *Dronc* functions in enteroblasts. The specific accumulation and activation of Dronc in EBs facilitates the cleavage of an unknown substrate “X” that either directly or indirectly could tune down Notch and Insulin pathways, thus preventing gut hyperplasia and the EB entry into EC differentiation program. (B) ISCs (white cell) are able to self-renew or differentiae into EB cells (Transition 1A) or into EE cells (Transition 1B). The EB (orange cell) can then differentiae into ECs (Transition 2, large grey cells). Dronc independently of effector caspases prevents the transition 2, thus inducing EB quiescence.

### Tumour suppressor role of caspases beyond apoptosis

The evasion of apoptosis is one of the hallmarks of tumour cells, and reasonably, the caspase regulation of apoptosis is one of the main tumour suppressor mechanism linked to these enzymes [55]. However, caspase activity in the intestinal system seems to be coupled to alternative tumour suppressor mechanisms. Along these lines, caspases can block the excess of proliferation and differentiation of intestinal stem cells [32, 56]. This clearly prevents gut hyperplasia as well as tumour prone conditions. Independently, our *Drosophila* findings indicate that caspases can limit the differentiation of EBs (Fig 5B). Importantly, these caspase effects are phenocopied by human *caspase*-9 intestinal precursors and are not linked to apoptosis. Instead they rely on the ability of caspases to maintain the pool of intestinal precursor cells in a quiescence status; our data and [33, 57]). Conversely, caspase deficiency can route proliferative precursors towards the differentiation pathway, thus adverting a tumorigenic situation. Collectively, this data supports the hypothesis that caspases can act as tumour suppressor genes through different and highly tissue specific mechanisms, which are often fully independent of apoptosis. Finally, they vividly illustrate some of the potential difficulties in implementing therapeutic approaches based on caspase-modulating molecules.

## MATERIALS AND METHODS

### Fly Husbandry

All fly strains used are described at www.flybase.bio.indiana.edu unless otherwise indicated. Primary *Drosophila* strains and crosses were routinely maintained on Oxford fly food. The fly food has three basic components that should be properly mixed (food base, Nipagin mix and Acid mix). The Exact amount of the three components are described next.

Base composition per litre of fly food:

agar (3 gr/l, Fisher Scientific UK Ltd, BP2641-1), malt (64.3 gr/l), molasses (18.8 gr/l), maize (64.3 gr/l), yeast (13 gr/l), soya (7.8 gr/l), water (1 l)
Nipagin mix per litre of fly food:

Methyl-4-hydroxybenzoate (nipagin) (2.68 gr, scientific labs, cat. W271004-10KG-K), Absolute ethanol (25.1 ml, Fisher), Water (1.3 ml)
Acid mix per litre of fly food:

5% phoshporic acid (Phosphoric Acid 85% Insect Cell Culture, cat. P5811-500G) in Propionic acid (5 ml, Propionic Acid Free Acid Insect Cell Culture, cat. P5561-1L)

Live yeast was added to each tube before transferring adult flies unless specified otherwise. Specific experiments also used Drosophila Quick Mix Medium Food (Blue) obtained from Blades Biological Ltd (DTS 070). 1g of Drosophila Quick Mix Medium (Blue) was mixed with 3mL of ddH_2_0 in these experiments.

### Conditional Knockout and Experimental regime during adulthood

Females were crossed with males at 18°C to prevent developmental lethality and transgene expression before adulthood. Adult female flies were then left mating with their siblings during 48h and finally transferred to 29 °C. At 29 °C the Gal80 repression of Gal4 is not effective allowing transgene expression. Until dissection, experimental specimens were transferred every two days to vials with fresh food containing yeast, or without yeast when using the Drosophila Quick Mix Media (Appendix Fig 1A).

### Paraquat treatment

The Drosophila were dry starved for 4 hours. Subsequently the flies were transferred to an empty fly vial in which the fly food was replaced by a flug soaked in a solution of 5% sucrose and 6.5mM Paraquat. Flies were left in this vial for 16 hours prior dissection.

### Full description of experimental genotypes

- *w*^1118^ DBS-S-QF, UAS-mCD8-GFP, QUAS-tomato-HA (Fig 1 A,D,G and Appendix Fig 1B)
- *w*^1118^; *esg*-Gal4 UAS-CD8-GFP / CYO; UAS-Histone-RFP TubG80^ts^ / TM6B (Fig 1 B,C,E,F,H,I and Appendix Fig 1C,D)
- *w*^1118^; *esg*-Gal4 UAS-CD8-GFP/ UAS-P35; UAS-Histone-RFP TubG80^ts^ / UAS-P35 (Fig1 G-I)
- *w*^1118^; *Dronc*^KO^ / *Dronc*^L29^ (Appendix Fig 2C)
- *w*^1118^; *Dronc*^KO^ / + (Appendix Fig 2D)
- *w*^1118^; *Dronc*^KO^ / *Dronc*^KO-DroncWT-Suntag-HA^ (Appendix Fig 2E)
- *w*^1118^; *esg*-Gal4 UAS-CD8-GFP / QUAS-LacZ; TubG80ts UAS-Histone-RFP *Dronc*^KO^ / UAS-Flp FRT *Dronc-GFP-APEX* FRT QF; (Appendix Fig 3A and B)
- *w*^1118^; *esg*-Gal4 UAS-CD8-GFP: TubG80^ts^ UAS-Histone-RFP *Dronc*^KO^ /SM6A-TM6B (Figure 2A,C-E; Appendix Fig. 3C-D, F-I, L-N and Appendix Fig 4 A and D)
- *w*^1118^; *esg*-Gal4 UAS-CD8-GFP / +; TubG80^ts^ UAS-Histone-RFP *Dronc*^KO^ / UAS-FLP FRT *Dronc-GFP-APEX* FRT Suntag-HA-Cherry (Fig. 2B-E, and Appendix Fig3 C-D and F-H)
- *w*^1118^; esg-Gal4 UAS-CD8-GFP / +; TubG80ts UAS-Histone-RFP *Dronc*^*KO*^ / UAS-FLP FRT *Dronc-GFP-APEX* FRT Dronc-WT-Suntag-HA-Cherry (Appendix Figure 3 E)
- *w*^1118^; *esg*-Gal4 UAS-CD8-GFP / +; TubG80^ts^ UAS-Histone-RFP *Dronc*^KO^ / UAS-FLP FRT *Dronc-GFP-APEX* FRT Dronc-FLCAEA-Suntag-HA-Cherry (Appendix Fig. 3J- and L-N)
- *w*^1118^; *esg*-Gal4 UAS-CD8-GFP / +; TubG80^ts^ UAS-Histone-RFP *Dronc*^KO^ / UAS-FLP FRT *Dronc-GFP-APEX* FRT Dronc-∆CAEA-Suntag-HA-Cherry (Appendix Fig. 3K and L-N)
- *w*^1118^; *esg*-Gal4 UAS-CD8-GFP / UAS-P35; TubG80ts UAS-Histone-RFP *Dronc*^KO^ / UAS-P35 (Appendix Fig. 4B and D)
- *w*^1118^; *esg*-Gal4 UAS-CD8-GFP / UAS-Drice RNAi UAS-Decay RNAi; TubG80ts UAS-Histone-RFP DroncKO12 / UAS-Damm RNAi UAS-DCP1 RNAi (Appendix Fig. 4C and D)
- *w*^1118^; + / QUAS-LacZ; *delta*-Gal4 UAS-GFP TubG80^ts^ UAS-Histone-RFP *Dronc*^KO^ / UAS-Flp FRT FRT *Dronc-GFP-APEX* FRT QF; (Appendix Fig. 3O and P)
- *w*^1118^; +/+; *delta*-Gal4 UAS-GFP TubG80^ts^ UAS-Histone-RFP *Dronc*^KO^/ TM6B (Fig. 2F,H-I and Appendix Fig 3Q)
- *w*^1118^; +/+; *delta*-Gal4 UAS-GFP TubG80^ts^ UAS-Histone-RFP *Dronc*^KO^ / UAS-FLP FRT FRT *Dronc-GFP-APEX* FRT Suntag-HA-Cherry (Fig. 2G-I and Appendix Fig 3Q)
- yw UAS-CD8-GFP/ *w*^1118^; +/+; *Dronc*-Gal4 / TM6B (Appendix Fig 5A)
- *w*^1118^; + / +; *Actin*-*Dronc*-GFP-MYC / TM6B (Fig. 3A; Appendix Fig 5C and D)
- w^1118^ / Su(H)-LacZ; + / +; Actin-*Dronc*-GFP-MYC / TM6B (Fig 3B)
- *w*^1118^ DBS-S-QF, UAS-mCD8-GFP, QUAS-tomato-HA / Su(H)-LacZ; +/+; +/+ (Fig. 3C and Appendix Fig 5E)
- *w*^1118^ UAS-Notch RNAi; *esg*-Gal4 UAS-CD8-GFP / +; TubG80ts UAS-Histone-RFP *Dronc*^KO^ / + (Fig. 4A and C)
- *w*^1118^ UAS-Notch RNAi; *esg*-Gal4 UAS-CD8-GFP / +; TubG80ts UAS-Histone-RFP *Dronc*^KO^ / UAS-FLP FRT *Dronc-GFP-APEX* FRT Suntag-HA-Cherry (Figure 4B and C)
- *w*^1118^; *esg*-Gal4 UAS-CD8-GFP / UAS-Notch Intra; TubG80ts UAS-Histone-RFP *Dronc*^KO^ / + (Fig. 4D and F)
- *w*^1118^; *esg*-Gal4 UAS-CD8-GFP / UAS-Notch Intra; TubG80ts UAS-Histone-RFP *Dronc*^KO^ / UAS-Flp FRT Dronc-GFP-APEX FRT QF (Fig. 4E and F)
- *w*^1118^; *esg*-Gal4 UAS-CD8-GFP / UAS-Insulin Receptor DN; TubG80ts UAS-Histone-RFP *Dronc*^KO^ / + (Fig. 4G and 4I)
- *w*^1118^; *esg*-Gal4 UAS-CD8-GFP / UAS-Insulin Receptor DN; TubG80ts UAS-Histone-RFP *Dronc*^KO^ / UAS-Flp FRT Dronc-GFP-APEX FRT QF (Fig. 4H and I)

### Molecular cloning and plasmid generation details

All PCRs were performed with Q5 High-Fidelity polymerase from New England Biolabs (NEB, M0492L). Standard subcloning protocols and HiFi DNA Assembly Cloning Kit (NEB, E5520S) were used to generate all the DNA plasmids. Transgenic lines expressing the new *Dronc* rescue constructs were obtained by attP/attB PhiC31-mediated integration. To this end, all the DNA plasmids were injected in *Drosophila* embryos containing the *Dronc*^KO^-reintegration site using Bestgene Inc. Fly strains generated will be deposited at the Bloomington Stock Centre. Whilst resource transfer is completed, reagents will be provided upon request. Description of RIV-Gal4 and plasmid backbones used for generating the different DNA rescue constructs can be found in Baena-Lopez et al 2013., [37].

### *pTV-Cherry-Dronc*^*KO*^. *Dronc* targeting vector

We first amplified the homology arms for generating the gene targeting of *Dronc* by PCR. Genomic DNA extracted through standard protocols from *w*^1118^ flies was used as a template. The sequences of the primers used are as follow:

5’ homology arm Forward primer with NotI restriction site:

5’ attatGCGGCCGCAAGTTGATGCAGCCTTCTGC 3’
5’ homology arm Reverse primer with KpnI restriction site:

5’ aattatGGTACCTCCGGTGACTCCGCTTATTGG 3’
3’ homology arm Forward primer with bglII restriction site:

5’ attgccaAGATCTACTGGACATTTTATCATTCC 3’
3’ homology arm Reverse primer with AvrII restriction site:

5’ attatggCCTAGGTCAAATCTGTTAATTTACG 3’

We then subcloned the PCR products into the targeting vector pTV-Cherry. The 5’ and 3’ homology arms were cloned into pTV-Cherry as a NotI-KpnI and BglII-AvrII fragments, respectively. The Dronc^KO^ behaves as previously described null alleles of the gene and is homozygous lethal during pupal development. The molecular validation of the allele was also carried out by PCR. First, we extracted the DNA from 10 larvae using the Quick genomic DNA prep protocol described in the link below. http://francois.schweisguth.free.fr/protocols/Quick_Fly_Genomic_DNA_prep.pdf.

The sequence of the primers used for performing the PCR were:

Forward KO validation primer:

5’ TGAGCAGCTGTTGGCATTAGG 3’
Reverse KO validation primer:

5’ AGAGAATACCAATCATACTGC 3’

The PCR conditions were 30 sec at 64 °C of annealing temperature and 1 min at 72 °C of extension. The PCRs was made using the Q5 High-Fidelity polymerase from New England Biolabs (NEB, M0492L).

### RIV-Dronc^*KO-Gal4*^

RIV-Gal4 plasmid was injected in the *Dronc*^*KO*^-reintegration site for generating the *Dronc*^*KO*-Gal4^ line.

### RIV-DroncKO-Dronc WT-Suntag-HA

Two fragments were amplified by PCR using the full-length wild-type cDNA of *Dronc* as a template (LP09975 cDNA clone from Drosophila Genomics Resource Centre). This cDNA also contained the flanking untranslated regions of *Dronc* in 5’ and 3’. We also appended a Suntag-HA-tag to the C-terminal end of the *Dronc* open reading frame, before the stop codon. The PCR fragments were then subcloned using HiFi DNA Assembly Cloning Kit into a PUC57 plasmid backbone previously opened with NotI-KpnI. The whole fragment was then transferred into the RIV(MCS-white[37]) as a NotI-KpnI fragment. The plasmid was finally injected in the *Dronc*^KO^-reintegration site. The primer sequences used for generating the rescue plasmid are next indicated.

Forward primer 1:

5’ tctagAGGGGGCTACCCATACGACGTCCCTGACTATGCGTAAgctagcttgccgccactggacattttatcattccgg 3’
Forward primer 2:

5’ gacggccagtgcggccgcagatctcctaggccggaacgcgtggaagccatatccggaatgcagccgccggagctcgagattgg 3’
Reverse primer 1:

5’ tctagAGGGGGCTACCCATACGACGTCCCTGACTATGCGTAAgctagcttgccgccactggacattttatcattccgg 3’
Reverse primer 2:

5’ caaaaattagactctttggccttagtcgggtaccggcgcgccactcgagcactagtTCTGGCTGTGTatatactgg 3’

Importantly, flies expressing this construct are viable, fertile and morphologically normal. Only the adult wings are less transparent than normal and sometimes do not expand normally.

### RIV-Dronc^*KO-FRT-DroncWT-GFP-Apex-FRT QF*^. Conditional Dronc allele followed by QF

We first modified the WT cDNA of *Dronc* adding a GFP-Apex2 chimeric fragment. This fragment was appended in frame before the stop codon of *Dronc* in order to facilitate biochemical approaches not used in this manuscript. The cDNA of *Dronc* and the GFP-Apex2 chimeric fragments were amplified by PCR from the plasmids *dronc*^*KO-Dronc-WT*^ and pCDNA-conexing-EGFP-APEX2 (addgene #49385), respectively. The sequences of the primers used to that end were:

Forward primer 1:

5’ tccggagcggccgcagatctCCGGAACGCGTGGAAGCCATATCCG 3’
Forward primer 2:

5’ TCCCGGGTTTTTCAACGAAggcggaatggtgagcaagggcgaggagc 3’
Reverse primer 1:

5’ tcgcccttgctcaccattccgccTTCGTTGAAAAACCCGGGATTG 3’
Reverse primer 2:

5’ atgtccagtggcggcaagctagcttaggcatcagcaaacccaagc 3’

The PCR fragments were then subcloned using HiFi DNA Assembly Cloning Kit into RIV-*Dronc*^*KO-Dronc-WT-Suntag-HA*^ previously opened with BglII-NheI. The entire construct was finally transferred into RIV (FRT-MCS2-FRT QF; *pax*-Cherry[37]) as a NotI-SpeI fragment. Notice that the extra sequences appended to *Dronc* (the FRT sites and the QF fragment) do not compromise the rescue ability of the construct. Indeed, these flies are identical to *RIV-Dronc*^*KO-Dronc-WT-Suntag-HA*^. Homozygous flies expressing QF upon FRT-rescue cassette excision die during metamorphosis indicating this allele in such configuration behaves as previously described null alleles.

### RIV-Dronc^*KO FRT-DroncWT-GFP-Apex-FRT Suntag-HA-Cherry*^. Conditional Dronc allele Suntag-HA-Cherry

We subcloned a newly designed PCR product Suntag-HA-Cherry in PUC57-Dronc^KO-Dronc WT-Suntag-HA^ by using HiFi DNA Assembly Cloning Kit. The PUC57-Dronc^KO-Dronc WT-Suntag-HA^ backbone vector was opened with AvrII-NsiI. The primers used for generating the Suntag-HA-Cherry peptide were:

Forward primer 1:

5’gccagtgcggccGCagatctCCTAGGcccgggtttttcaacgaagggggcgaggagttgctgaGCAAAAATTATCATTTG GAGAacgaagtagcacgactaaag 3’
Forward primer 2:

5’gtagcacgactaaagaaagggtccggatcgggttctagagggggctacccatacgacgtccctgactatgcgGGGaattCCAACatggtgagcaaggg cg 3’
Reverse primer 1:

5’cgcccttgctcaccatGTTGGaattCCCcgcatagtcagggacgtcgtatgggtagccccctctagaacccgatccggaccctttctttagtcgtgctac 3’
Reverse primer 2:

5’ggacgagctgtacaagtaagacgtcgtcgaGGGTACCTctagcttgccgccactggacattttatcattccggatgcatttttaaccgcatttatgt 3’

Then the Suntag-HA-Cherry was extracted from the new PUC*57-Dronc*^*KO-Dronc-WT-Suntag-HA-Cherry*^ vector as a NotI-ClaI fragment and transferred to *RIV-Dronc*^*KO FRT-DroncWT-GFP-Apex-FRT QF*^ previously opened with AvrII--ClaI. Before ligation, NotI and AvrII sites were blunted. Homozygous flies expressing Suntag-HA-Cherry peptide under the physiological regulation of *Dronc* die during metamorphosis indicating this allele behaves as previously described null alleles.

### RIV-Dronc^*KO FRT-DroncWT-GFP-Apex-FRT Dronc-FLCAEA-Suntag-HA-Cherry*^. Conditional Dronc allele FL-CAEA-Suntag-HA-Cherry

We first generated two point mutations through gene synthesis (Genewizz) in the wild-type cDNA of *Dronc* that caused the following amino acid substitutions; C318A and E352A. This version of *Dronc* is enzymatically inactive (C318A), and cannot be either processed during the proteolytic activation steps of *Dronc* (E352A). This fragment was subcloned in PUC*57-Dronc*^*KO-Dronc-WT-Suntag-HA-Cherry*^ as a BglII-XmaI fragment, thus replacing the wildtype version of *Dronc* by the mutated. Finally, the DNA sequence was transferred to the *RIV-Dronc*^*KO FRT-DroncWT-GFP-Apex-FRT QF*^ plasmid as an AvrII-ClaI fragment. Homozygous flies expressing this mutant form of *Dronc* die during metamorphosis indicating this allele behaves as previously described null alleles.

### RIV-Dronc^*KO FRT-DroncWT-GFP-Apex-FRT Dronc-deltaCAEA-Suntag-HA-Cherry*^. Conditional Dronc allele delta-CAEA-Suntag-HA-Cherry

We generated a PCR product that deletes the CARD domain of Dronc using the following primers and as template for the PCR *RIV-Dronc*^*KO FRT-DroncWT-GFP-Apex-FRT Dronc-FLCAEA-Suntag-HA-Cherry*^.

Forward primer 1:

5’ ggccagtgcggccGCCCTAGGGTTTaaacggggaatgggcaattGtctggatgcggcc 3’
Reverse primer 1:

5’ catGTTGGaattccccgcatagtcagggacgtcgtatgggtagccccc 3’

The PCR product was subcloned in PUC*57-Dronc*^*KO-Dronc-Suntag-HA-Cherry*^ as a NotI-EcoRI fragment, thus inserting the truncated and catalytically inactive version of Dronc in frame with the Suntag-HA-Cherry peptide. Finally, the DNA sequence was transferred to the *RIV-Dronc*^*KO FRT-DroncWT-GFP-Apex-FRT QF*^ plasmid as an AvrII-PasI fragment. Homozygous flies expressing this mutant form of *Dronc* die during metamorphosis indicating this allele behaves as previously described null alleles.

### RIV-DroncKO FRT-DroncWT-GFP-Apex-FRT Dronc-WT-Suntag-HA-Cherry. Conditional Dronc allele Dronc-WT-Suntag-HA-Cherry

We generated a PCR product using as a template for the PCR *RIV-Dronc*^*KO-Dronc WT-Suntag-HA*^ and using the following primers.

Forward primer 1:

5’ ggccagtgcggccgcagatctcctaggccggaacgcgtggaagccatatccggaatgcagccgccgga 3’
Reverse primer 1:

5’ cccttgctcaccatGTTGGaattCCCcgcatagtcagggacgtcgtatggg 3’

The PCR product was subcloned in PUC*57-Dronc*^*KO-Dronc-Suntag-HA-Cherry*^ as a NotI-EcoRI fragment, thus inserting in frame the wild type version of Dronc in frame with the Suntag-HA-Cherry peptide. Finally, the DNA sequence was transferred to the *RIV-Dronc*^*KO FRT-DroncWT-GFP-Apex-FRT QF*^ plasmid as an AvrII-PasI fragment. Heterozygous flies expressing this mutant form rescue the pupal lethality associated with *Dronc* insufficiency.

### Plasmid generation details of Actin-Dronc-CA-GFP-Myc

We first generated one-point mutation through gene synthesis (Genewizz) in the wild-type cDNA of *Dronc* that causes the following amino acid substitution C318A. This version of *Dronc* is enzymatically inactive. In addition we appended a Suntag and a HA peptide sequence to the C-terminus, in frame with the open reading frame (ORF) of Dronc. Downstream of the ORF we included the 3’UTR of *Dronc* present in the genomic locus. Extra restriction sites were added at the 5’ and 3’ ends of the construct to facilitate future subcloning projects. The entire construct was subcloned in PUC*57* as a Not-KpnI fragment. We then opened this vector with SmaI and NheI; this enzymatic digestion eliminates the C-terminal tagging of Dronc (Suntag-HA) whilst retaining the 3’UTR. Using HiFi DNA assembly, we inserted a PCR product that encodes for a modified version of GFP with a Myc tag appended at the C-terminal end. The primers used to amplify the modified GFP-Myc were:

Forward primer 1:

5’ GCTTTAATAAGAAACTCTACTTCAATcccgggtttttcaacgaagggggcATGATCAAGATCGCCACCAGGAAGTACC 3’
Forward primer 2:

5’ CGTGACCGCCGCCGGGATCACGGAAACCGATGGCGAGCTGTTCACCGGGGTGG 3’
Reverse primer 1:

5’ CCACCCCGGTGAACAGCTCGCCATCGGTTTCCGTGATCCCGGCGGCGGTCACG 3’
Reverse primer 2:

5’gataaaatgtccagtggcggcaagctagCttacaggtcctcctcgctgatcagcttctgctcGTTAGGCAGGTTGTCCACCCTCATCAGG 3’

The template used to obtain the GFP-Myc PCR product was extracted from genomic DNA of flies containing the construct UAS-GC3Ai [58]. The construct was finally subcloned as a NotI-XhoI fragment in an Actin-polyA vector of the lab previously opened with NotI-PspXI. Sequence of the plasmid will be provided upon request until the vector is deposited in a public repository.

### Immunohistochemistry

Adult mated female *Drosophila* Intestines were dissected in ice-cold PBS. Following dissection, the intestines were fixed by immersing for 6 seconds in wash solution (0.7% NaCl, 0.05% Triton X-100) heated to approximately 90°C. Subsequently the intestines were rapidly cooled in Ice-cold wash solution. The intestines were then rapidly washed in PBT (0.3%) before blocking for at least 1 hour in 1% BSA-PBT (0.3%). Primary antibodies were incubated overnight at 4°C and secondary antibodies at room temperature for two hours, diluted in blocking solution. Primary antibodies used were: Goat Anti-GFP (1:200, Abcam, ab6673) Chicken anti-Beta-galactosidase (1:200, Abcam, Ab9361); Rabbit Anti-HA (1:1000, Cell Signalling, C29F4); Rabbit anti-Pdm1 (1:2000, kind gift from Yu Cai), Mouse Anti-Armadillo (1:50, DSHB, N2 7A1 ARMADILLO-c); Mouse Anti-Prospero (1:20; DSHB, MR1A), Rabbit Anti-P35 (1:100, Novus Biologicals, NB100-56153) The secondary antibodies used were: DAPI (1:1000, Thermo Scientific, 62248); Goat Alexa 488 anti-Chicken (1:200, Life technologies, A11039); Donkey Alexa-488 anti-Goat (1:200, Life Technologies, A1105); Donkey Alexa-555 anti-Rabbit (1:200, Life Technologies, A31572),Donkey Alexa-647 anti-Rabbit (1:200, Life Technologies, A31573); Donkey Alexa-555 Anti-Mouse (1:200, Life Technologies, A31570) & Donkey Alexa-647 Anti-chicken (1:200, Jackson, 703-605-155)

### Imaging of fixed samples

Fluorescent imaging of the R5 posterior region [59] of *Drosophila* intestines were performed using the Olympus Fluoview FV1200 and associated software. Z-stacks were taken using either the 40x or 10x lenses. Acquired images were processed and quantified using automated Fiji/ImageJ [60, 61] macros or manually when automatisation was not possible. Generally, Z-stacks were projected, and the channels split. The “despeckle” tool was utilised to remove noise. The image was then “thresholded” and the “watershed” tool used to segment joined cells. To count the number of objects, the “Analyse Particles…” function was utilised in automated quantification, and for manual counting, “cell counter”. Figures were produced with Adobe Illustrator CC 2017.

### Statistical Analysis

Microsoft Excel was used to complete the basic numerical preparation of the data. The data was subsequently collated in Graph Pad Prism (8). The “identity outliers” analysis was utilised using the ROUT method with a Q value of 1% (All N numbers listed in figures are prior to this analysis). The cleaned data was then tested for normality using the D’Agostino-Pearson omnibus normality test except for qPCR data which was tested using the Shapiro-Wilk normality test. All subsequent analysis is referenced in Figure Legends. The P value format used is as follows: ns = P > 0.05, * = P ≤ 0.05, ** = P ≤ 0.01, *** = P ≤ 0.001 and **** = P ≤ 0.0001.

### RNA Extraction and cDNA Synthesis of Drosophila Intestines

Adult mated female *Drosophila* Intestines were dissected in ice-cold PBS. Following dissection, the intestines were transferred to autoclaved Eppendorf tubes containing 350μL of RLT Buffer plus from the RNeasy Plus Micro Kit (QIAGEN Cat. No 74034) with 1% v/v of 2-mercaptoethanol (Sigma-Aldrich M6250). The intestines were homogenised using a new 1.5ml pestle (Kimble 749521-1590) for each sample. The RNA was then extracted using the protocol and materials outlined in the RNeasy Plus Micro Kit. RNA quantity in samples were assessed twice using the Nanodrop Lite Spectrophotometer (Thermo Fisher Scientific) and the average determined. 500μg of RNA was synthesized to cDNA (Thermo Fisher Scientific Maxima First Strand cDNA Synthesis Kit – K1671). QPCR was performed using reagents and protocols from QIAGEN QuantiTect SYBR^⊙^ Green PCR Kit (Cat No./ID: 204145) and using the QIAGEN Rotor-Gene Q.

The Primer sequences were:

Rpl32 [62]:

Forward: 5’ ATGCTAAGCTGTCGCACAAATG 3’
Reverse: 5’ GTTCGATCCGTAACCGATGT 3’
Alpha-Trypsin* (PD44223):

Forward: 5’ ATGGTCAACGACATCGCTGT 3’
Reverse: 5’ CTGGCTCTGGCTAACGATGT 3’
Amylase-D* (PD40005):

Forward: 5’ GCATAGTGTGCCTCTCCCTC 3’
Reverse: 5’ TACGACCGGATGCGTAGTTG 3’
Jon65Aiii [63]:

Forward: 5’ AACACCTGGGTTCTCACTGC 3’
Reverse: 5’ TCAGGGAAATGTCGTTCCTC 3’

* Primers were sourced from the QPCR FlyPrimerBank tool [64].

## ACKNOWLEDGMENTS

The authors would like to thank Irene Miguel-Aliaga (Imperial College London) for sharing various flies and reagents. Yu CAI (Temasek Life Sciences Laboratory Limited) for generously sharing the Pdm1 antibody. Joaquín Navascués for generously sharing reagents and protocols. Antonello Zeus for his intellectual input at the initial stages of the project. The Caspase Lab members for their critical reading of the manuscript and invaluable suggestions. This work has been supported by Cancer Research UK 560C49979/A17516 and the John Fell Fund from the University of Oxford 162/001. L.A.Baena-Lopez is a CRUK Career Development Fellow (C49979/A17516) and an Oriel College Hayward Fellow. J. Alonso was a CRUK postdoctoral scientist associated to the grant code previously described. D. Antoni. Nahotko. was a summer student supported by the ERASMUS (+) program. L. Arthurton is a PhD student supported by the Edward Penley Abraham Research Fund.

## CONTRIBUTIONS

L.A.B-L. was responsible for the initial conceptualisation of the work. D. N., J. A, and L.A.B-L were responsible for the plasmid generation and molecular biology protocols and the stock preparation of the conditional alleles. The majority of the experimental design and discussion was elaborated by L. A. and L.A.B-L. The experimental work was mostly performed by L. A. The manuscript was written and corrected by L. A. and L.A.B.-L. The figures of the manuscript were produced by L. A. and L.A.B-L. All co-authors commented on the manuscript and approve the submission.

## REFERENCES

1. Xu, D.C., L. Arthurton, and L.A. Baena-Lopez, Learning on the Fly: The Interplay between Caspases and Cancer. Biomed Res Int, 2018. 2018: p. 5473180.

2. Baena-Lopez, L.A., et al., Non-apoptotic Caspase regulation of stem cell properties. Semin Cell Dev Biol, 2018. 82: p. 118–126.

3. Miles, W.O., N.J. Dyson, and J.A. Walker, Modeling tumor invasion and metastasis in Drosophila. Dis Model Mech, 2011. 4(6): p. 753–61.

4. Miguel-Aliaga, I., H. Jasper, and B. Lemaitre, Anatomy and Physiology of the Digestive Tract of Drosophila melanogaster. Genetics, 2018. 210(2): p. 357–396.

5. Crawford, E.D. and J.A. Wells, Caspase substrates and cellular remodeling. Annu Rev Biochem, 2011. 80: p. 1055–87.

6. Parrish, A.B., C.D. Freel, and S. Kornbluth, Cellular mechanisms controlling caspase activation and function. Cold Spring Harb Perspect Biol, 2013. 5(6).

7. Goyal, L., et al., Induction of apoptosis by Drosophila reaper, hid and grim through inhibition of IAP function. EMBO J, 2000. 19(4): p. 589–97.

8. Srinivasula, S.M., et al., sickle, a novel Drosophila death gene in the reaper/hid/grim region, encodes an IAP-inhibitory protein. Curr Biol, 2002. 12(2): p. 125–30.

9. Wing, J.P., et al., Drosophila sickle is a novel grim-reaper cell death activator. Curr Biol, 2002. 12(2): p. 131–5.

10. Christich, A., et al., The damage-responsive Drosophila gene sickle encodes a novel IAP binding protein similar to but distinct from reaper, grim, and hid. Curr Biol, 2002. 12(2): p. 137–40.

11. Steller, H., Regulation of apoptosis in Drosophila. Cell Death Differ, 2008. 15(7): p. 1132–8.

12. Silke, J. and P. Meier, Inhibitor of apoptosis (IAP) proteins-modulators of cell death and inflammation. Cold Spring Harb Perspect Biol, 2013. 5(2).

13. Huh, J.R., et al., The Drosophila inhibitor of apoptosis (IAP) DIAP2 is dispensable for cell survival, required for the innate immune response to gram-negative bacterial infection, and can be negatively regulated by the reaper/hid/grim family of IAP-binding apoptosis inducers. J Biol Chem, 2007. 282(3): p. 2056–68.

14. Leulier, F., et al., The Drosophila inhibitor of apoptosis protein DIAP2 functions in innate immunity and is essential to resist gram-negative bacterial infection. Mol Cell Biol, 2006. 26(21): p. 7821–31.

15. Muro, I., K. Monser, and R.J. Clem, Mechanism of Dronc activation in Drosophila cells. J Cell Sci, 2004. 117(Pt 21): p. 5035–41.

16. Rodriguez, A., et al., Dark is a Drosophila homologue of Apaf-1/CED-4 and functions in an evolutionarily conserved death pathway. Nat Cell Biol, 1999. 1(5): p. 272–9.

17. Xu, D., et al., The CARD-carrying caspase Dronc is essential for most, but not all, developmental cell death in Drosophila. Development, 2005. 132(9): p. 2125–34.

18. Cheng, T.C., et al., A Near-Atomic Structure of the Dark Apoptosome Provides Insight into Assembly and Activation. Structure, 2017. 25(1): p. 40–52.

19. Baena-Lopez, L.A., et al., Novel initiator caspase reporters uncover previously unknown features of caspase-activating cells. Development, 2018. 145(23).

20. Ohlstein, B. and A. Spradling, The adult Drosophila posterior midgut is maintained by pluripotent stem cells. Nature, 2006. 439(7075): p. 470–4.

21. Micchelli, C.A. and N. Perrimon, Evidence that stem cells reside in the adult Drosophila midgut epithelium. Nature, 2006. 439(7075): p. 475–9.

22. Jiang, H. and B.A. Edgar, Intestinal stem cells in the adult Drosophila midgut. Exp Cell Res, 2011. 317(19): p. 2780–8.

23. Li, H. and H. Jasper, Gastrointestinal stem cells in health and disease: from flies to humans. Dis Model Mech, 2016. 9(5): p. 487–99.

24. Zacharioudaki, E. and S.J. Bray, Tools and methods for studying Notch signaling in Drosophila melanogaster. Methods, 2014. 68(1): p. 173–82.

25. Salazar, J.L. and S. Yamamoto, Integration of Drosophila and Human Genetics to Understand Notch Signaling Related Diseases. Adv Exp Med Biol, 2018. 1066: p. 141–185.

26. Lieber, T., et al., Antineurogenic phenotypes induced by truncated Notch proteins indicate a role in signal transduction and may point to a novel function for Notch in nuclei. Genes Dev, 1993. 7(10): p. 1949–65.

27. Wang, H., et al., The role of Notch receptors in transcriptional regulation. J Cell Physiol, 2015. 230(5): p. 982–8.

28. Hakim, R.S., K. Baldwin, and G. Smagghe, Regulation of midgut growth, development, and metamorphosis. Annu Rev Entomol, 2010. 55: p. 593–608.

29. Kapuria, S., et al., Notch-mediated suppression of TSC2 expression regulates cell differentiation in the Drosophila intestinal stem cell lineage. PLoS Genet, 2012. 8(11): p. e1003045.

30. Quan, Z., et al., TSC1/2 regulates intestinal stem cell maintenance and lineage differentiation through Rheb-TORC1-S6K but independently of nutritional status or Notch regulation. J Cell Sci, 2013. 126(Pt 17): p. 3884–92.

31. Zhai, Z., J.P. Boquete, and B. Lemaitre, A genetic framework controlling the differentiation of intestinal stem cells during regeneration in Drosophila. PLoS Genet, 2017. 13(6): p. e1006854.

32. Jin, Y., et al., Brahma is essential for Drosophila intestinal stem cell proliferation and regulated by Hippo signaling. Elife, 2013. 2: p. e00999.

33. Xu, D., et al., Apoptotic block in colon cancer cells may be rectified by lentivirus mediated overexpression of caspase-9. Acta Gastroenterol Belg, 2013. 76(4): p. 372–80.

34. Antonello, Z.A., et al., Robust intestinal homeostasis relies on cellular plasticity in enteroblasts mediated by miR-8-Escargot switch. EMBO J, 2015. 34(15): p. 2025–41.

35. Ali, S., et al., Paraquat induced DNA damage by reactive oxygen species. Biochem Mol Biol Int, 1996. 39(1): p. 63–7.

36. Hay, B.A., T. Wolff, and G.M. Rubin, Expression of baculovirus P35 prevents cell death in Drosophila. Development, 1994. 120(8): p. 2121–9.

37. Baena-Lopez, L.A., et al., Accelerated homologous recombination and subsequent genome modification in Drosophila. Development, 2013. 140(23): p. 4818–25.

38. Chew, S.K., et al., The apical caspase dronc governs programmed and unprogrammed cell death in Drosophila. Dev Cell, 2004. 7(6): p. 897–907.

39. Doupe, D.P., et al., Drosophila intestinal stem and progenitor cells are major sources and regulators of homeostatic niche signals. Proc Natl Acad Sci U S A, 2018. 115(48): p. 12218–12223.

40. Ouyang, Y., et al., Dronc caspase exerts a non-apoptotic function to restrain phospho-Numb-induced ectopic neuroblast formation in Drosophila. Development, 2011. 138(11): p. 2185–96.

41. Leulier, F., et al., Systematic in vivo RNAi analysis of putative components of the Drosophila cell death machinery. Cell Death Differ, 2006. 13(10): p. 1663–74.

42. Xu, D., et al., The effector caspases drICE and dcp-1 have partially overlapping functions in the apoptotic pathway in Drosophila. Cell Death Differ, 2006. 13(10): p. 1697–706.

43. Zeng, X., C. Chauhan, and S.X. Hou, Characterization of midgut stem cell- and enteroblast-specific Gal4 lines in drosophila. Genesis, 2010. 48(10): p. 607–11.

44. Ryoo, H.D., T. Gorenc, and H. Steller, Apoptotic cells can induce compensatory cell proliferation through the JNK and the Wingless signaling pathways. Dev Cell, 2004. 7(4): p. 491–501.

45. Kamber Kaya, H.E., et al., An inhibitory mono-ubiquitylation of the Drosophila initiator caspase Dronc functions in both apoptotic and non-apoptotic pathways. PLoS Genet, 2017. 13(2): p. e1006438.

46. Amcheslavsky, A., et al., Plasma Membrane Localization of Apoptotic Caspases for Non-apoptotic Functions. Dev Cell, 2018. 45(4): p. 450–464 e3.

47. Guo, Z. and B. Ohlstein, Stem cell regulation. Bidirectional Notch signaling regulates Drosophila intestinal stem cell multipotency. Science, 2015. 350(6263).

48. Koch, U., R. Lehal, and F. Radtke, Stem cells living with a Notch. Development, 2013. 140(4): p. 689–704.

49. Ohlstein, B. and A. Spradling, Multipotent Drosophila intestinal stem cells specify daughter cell fates by differential notch signaling. Science, 2007. 315(5814): p. 988–92.

50. Amcheslavsky, A., J. Jiang, and Y.T. Ip, Tissue damage-induced intestinal stem cell division in Drosophila. Cell Stem Cell, 2009. 4(1): p. 49–61.

51. Jin, Y., et al., Intestinal Stem Cell Pool Regulation in Drosophila. Stem Cell Reports, 2017. 8(6): p. 1479–1487.

52. Reiff, T., et al., The Notch and EGFR signaling regulate caspase inhibitor Diap1 to match supply with intestinal demand. 2018: p. 493528.

53. Aram, L., K. Yacobi-Sharon, and E. Arama, CDPs: caspase-dependent non-lethal cellular processes. Cell Death Differ, 2017. 24(8): p. 1307–1310.

54. Bell, R.A.V. and L.A. Megeney, Evolution of caspase-mediated cell death and differentiation: twins separated at birth. Cell Death Differ, 2017. 24(8): p. 1359–1368.

55. Hanahan, D. and R.A. Weinberg, Hallmarks of cancer: the next generation. Cell, 2011. 144(5): p. 646–74.

56. Lee, D.J., et al., Regulation and Function of the Caspase-1 in an Inflammatory Microenvironment. J Invest Dermatol, 2015. 135(8): p. 2012–2020.

57. Asadi, M., et al., Expression Level of Caspase Genes in Colorectal Cancer. Asian Pac J Cancer Prev, 2018. 19(5): p. 1277–1280.

58. Schott, S., et al., A fluorescent toolkit for spatiotemporal tracking of apoptotic cells in living Drosophila tissues. Development, 2017. 144(20): p. 3840–3846.

59. Dutta, D., et al., Regional Cell-Specific Transcriptome Mapping Reveals Regulatory Complexity in the Adult Drosophila Midgut. Cell Rep, 2015. 12(2): p. 346–58.

60. Schindelin, J., et al., Fiji: an open-source platform for biological-image analysis. Nat Methods, 2012. 9(7): p. 676–82.

61. Rueden, C.T., et al., ImageJ2: ImageJ for the next generation of scientific image data. BMC Bioinformatics, 2017. 18(1): p. 529.

62. Gomez-Lamarca, M.J., et al., Activation of the Notch Signaling Pathway In Vivo Elicits Changes in CSL Nuclear Dynamics. Dev Cell, 2018. 44(5): p. 611–623 e7.

63. Bozler, J., et al., A systems level approach to temporal expression dynamics in Drosophila reveals clusters of long term memory genes. PLoS Genet, 2017. 13(10): p. e1007054.

64. Hu, Y., et al., FlyPrimerBank: an online database for Drosophila melanogaster gene expression analysis and knockdown evaluation of RNAi reagents. G3 (Bethesda), 2013. 3(9): p. 1607–16.

